# Polymer brush bilayer under stationary shear: A joint DFT, scaling theory and MD

**DOI:** 10.1101/2022.08.16.504146

**Authors:** Mike John Edwards (Majid Farzin)

## Abstract

The problem of polymer brush bilayer under stationary shear is studied by using the DFT, the scaling theory and MD simulations. Both theory and simulations confirm that the shear stress follows the universal power law 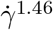 for the brush bilayers with interpenetration and in the absence of the interpenetration, the shear stress scales linearly with the shear rate. It is also revealed that the presence of explicit solvent molecules prevents the brushes to form and interpenetration zone and therefore with explicit solvents the shear stress scales linearly with shear rate. Therefore, this study strongly confirms that there is no sublinear regime in the world of polymer brush bilayer, neither by solvents nor by hydrodynamic effects. As long as there is an interpenetration zone, the superlinear regime dominates and in the absence of the interpenetration zone the linear regime dominates. Therefore, polymer brushes are not a good candidate for lubrication and all works suggesting that this system is a super lubricant are completely wrong.

## Introduction

Polymers are linear macromolecular structures that are already known as building blocks of life. They are composed of repeating units of atoms or molecules. Each repeating unit of polymers are known as monomers. The connectivity throughout monomers is established via covalent bonds in which the valence electrons of atoms or molecules are shared together. The covalent bonds are among the strongest bonds in nature. The Brownian motion of monomers which arise from collisions by interstitial water molecules, causes monomers to fluctuate. The fluctuations of monomers causes the whole chain to undergo all possible configurations in the long time measurement. The average chain extension plus other time-averaged quantities are key properties of polymers that physicists seek. The end-to-end distance *R*_*e*_ and the radius of gyration *R*_*g*_ ^1,6,20^ are two candidates for average chain extension, however, *R*_*g*_ is more accurate. Simple calculations show that 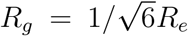. For point-like monomers with no internal structure, *R*_*e*_ = *aN* ^1*/*2^ that is called freely-jointed-chain model (FJC). Where, *N* is the number of monomers i.e. degree of polymerization and *a* is the monomer size or the *Kuhn* length. In the FJC model, the monomers distribute by the Gaussian distribution. If we assume that monomers have internal structure, the monomers will interact via excluded volume interactions. All polymer Physics approaches including perturbation expansion theory (PET), renormalization group theory (RG), scaling theory (ST), mean field theory (MFT) and density functional theory (DFT) reveal that 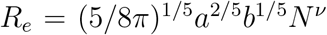. Where, *b* here denotes the second Virial coefficient and beers information about interactions between monomers.^1,6,20^ The Flory exponent *ν* is equal to 3*/*5 or more precisely 0.588.^1,6,20^ In essence, 1*/ν* is the fractal dimension of the linear polymer chains.^3^ As the excluded volume interactions between monomers turns on, distribution of monomers gets parabolic. ^1,6,20^ One of most interesting fact is that the average chain extension is a very sensitive function of molecular parameters, interactions between monomers and between monomers and solvent molecules. For instance, when the chain is suspended in a good solvent like water, *b* = 2.09 *a*^3^ so the chain swells, however, when it is suspended in a *θ* solvent *b* = 0 and the Gaussian model holds. ^1,6,20^ On the other hand, a polymer chain suspended in a poor solvent like Alcohol, feels *b <* 0 and chain collapses i.e. *R*_*e*_ *∼ N* ^1*/*3^.^1,6,20^

One of the most applicable forms of polymers is the polymer solution in which the linear chains are grafted to a flat surface by one end. This system is known as polymer brushes and has been the subject of a huge number of researches in the last decades. In this system, the steric repulsion among the monomers stretch the chains drastically in the perpendicular direction. The density functional theory (DFT) reveals that the average perpendicular chain extension in perpendicular direction or brush height is *R*_*n*_ = (*a*^2^*bσ*)^1*/*3^*N* and the mean lateral chain extension is the same as single isolated chain. ^2,19^ Polymer brush coated surfaces can have self healing properties and many others such as Glycol on the outside of the cell membrane and they play a key role in cells movement and interactions. ^2^

Nevertheless, here, I am interested in lubrication properties of two opposing brush covered surfaces that are located in the distance where the top and bottom chains intermediately interpenetrate. This system is known as polymer brush bilayer (PBB).^2,23–25^ The lubrication properties of the PBBs play a key role in the joints of humans and other mammals. The presence of aggregans in the synovial fluid of the mammalian joints are a good example. Recently, there has been a tough competition on establishing a correct theory for the equation of state of the PBBs in equilibrium and under stationary shear.^5,8–15^ Many experiments has been also done to test lubrication properties of the PBBs.^16–18^

In the coming two sections, I briefly describe the theoretical and simulation parts and in the third section, I show the results. At the final section, normally I discuss the results and conclude the research.

### Low shear rate scenario

One of the useful methods to study mechanics of a many body system is the DFT. In the DFT, the Euler-Lagrange equation of the system is calculated from a grand potential functional which is primarily based on a guess. There are also other methods such as Newton’s equation of motion and Hamilton-Jacobi method. In 1988, Hirz in his Master’s thesis applied the DFT method to polymer brush problem and obtained density profiles as well as brush height.^7^ Following the same procedure that Hirz did, the PBBs grand potential functional could be given as follows,

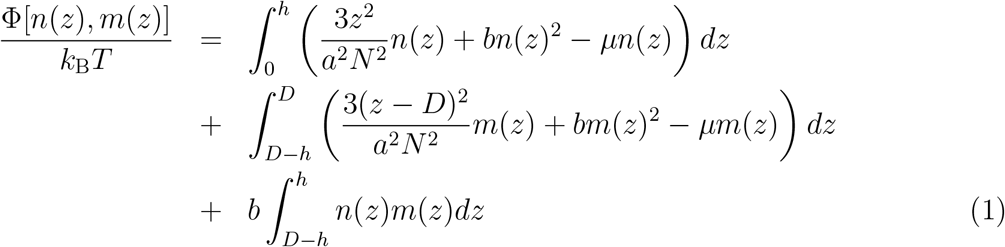

where *n*(*z*) and *m*(*z*) are the density profiles of the bottom and up brushes respectively, *h* the brush height, *μ* the chemical potential, *σ* the grafting density and *D* the distance between flat surfaces. Three Euler-Lagrange equations could be obtained plus one constraint which all are given as follows,

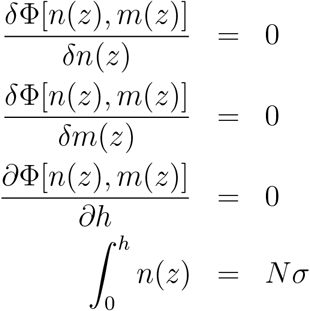

The first three equations say that equilibrium density profiles and height is calculated by setting functional derivatives of grand potential with respect those quantities to zero. The last equation says that the total number of particles are fixed. To obtain equilibrium *n*(*z*), *m*(*z*), *h* and *μ* that are primarily unknown quantities, the four equations above need to be solved simultaneously. The solutions of the above set of equations are given as follows,

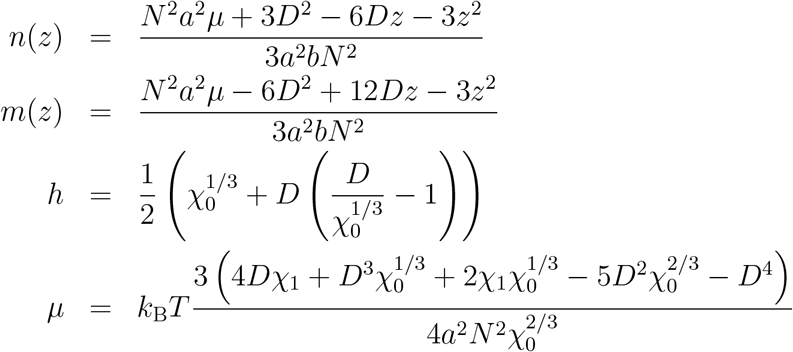

where *χ*_0_ and *χ*_1_ are volumes defined as follows,

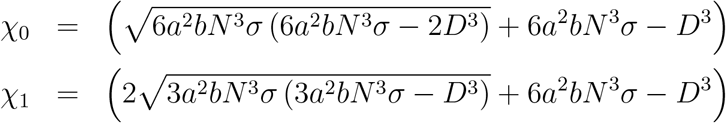

By plotting *h* as a function of each parameter, it turns out that *h* follows the following similarity relation,

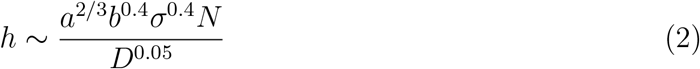

To calculate the equation of state of the PBBs, all calculated quantities put back into Eq. (1), and calculate the pressure via the Gibbs-Dohem relation Φ = *−PV* . The equation of state of the PBBs is calculated as follows,

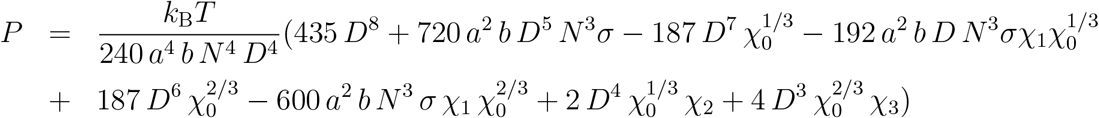

where the following volumes are defined,

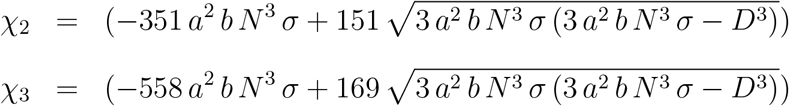

By plotting the pressure equation, it could be simply inferred that the pressure follows the following similarity relation,

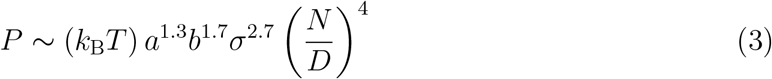

It turns out that, the equation of state scales as (*N/D*)^4^ which is very important outcome and violates all previous calculations. It reveals that pressure is very strongly sensitive to *N* and *D* unlike single brush. The quadratic dependence of PBBs pressure on the degree of polymerization as well as distance between surfaces originates from complex interactions between monomers.

The phenomenological arguments approaches a physical system by means of simple assumeptions and obtains the general behavior of that system. The scaling theory is based on the phenomenological arguments and is capable of solving complex physical problems simply. The problem of the PBB under shear seems to be very complicated at first sight. However, by using the following argument, it get easier to solve. Upon shearing the PBB, above a critical shear rate, the chains start stretching in the shear direction. This is similar to the equilibrium condition but with chains stretched laterally. We can use this concept to build our scaling theory. I introducw a scaling function in form of 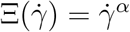 with 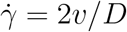 the shear rate and *v* the relative velocity of surfaces. The goal is to calculate the exponent *α*. From DFT calculations, we know the physical quantities at low shear rates. The normal stress is the same as Eq. (3) and the shear stress is obtained by multiplying the viscosity by the shear rate. To calculate the viscosity, we need to look deep inside the system in the molecular scale. Each particle in the system, travels freely an amount of time *τ* between two subsequent collisions with another particle. The corresponding distance is called mean-free-path *λ*. From statistical Physics, it is known that for each system, 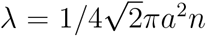 with *n* density of the system. The average density of the PBBs is *n* = 2*Nσ/D* so the mean-free-path becomes,

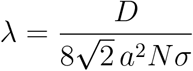

Again from statistical Physics, it is known that for each system *τ* = *λ/v*_*rms*_ with *v*_*rms*_ the root-mean-squared velocity of particles. For polymer chains, it is a bit tricky to calculate the root-mean-squared velocity. Since the monomers undergo Brownian motion, they perform diffusive movement rather than ballistic movement. To this reason, for polymer chains *v*_*rms*_ = *λ/D*_*G*_ with *D*_*G*_ the diffusion constant of the center of mass monomer. This literally means that every monomer diffuses the same as the center of mass monomer *D*_*G*_. In the next subsection, I replace the appropriate *D*_*G*_ to calculate the *λ* and I obtain the corresponding *τ* . When the correct form of the *τ* is known for each type of system (here, I mean the system with or without the HIs), the viscosity is obtained as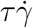

In the absence of HIs, the polymer dynamics is governed by Rouse dynamics.^20^ In this regime, when the shear rate exceeds a critical value, the shear forces dominate the fluctuation forces as well as the steric forces, and consequently, the chains start stretching in the shear direction. In the Rouse’s dynamic, the critical shear rate is equal to inverse of *τ*_*c*_ = *ξ a*^2^ *N* ^2*ν*+1^*/*3*π*^2^ *k*_B_*T* . Note that, this is the very longest Rouse’s relaxation time. For simplicity, it is useful to work with a dimensionless quantity 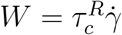 instead of shear rate. Now, step by step, I calculate the physical quantities both in linear and non-linear response regimes. The chain extensions at linear response regime are given as follows,

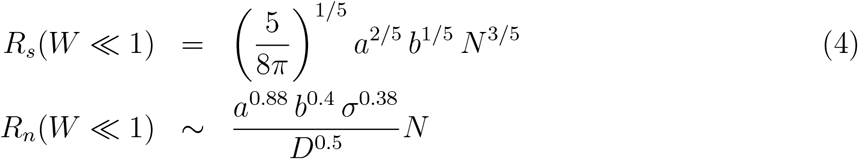

The diffusion constant of center of mass monomer in the Rouse’s dynamics with excluded volume interactions is given as *D*_*G*_ = *k*_B_*T/Nξ*.^20^ Therefore, the mean-free-time in this condition is given as follows,

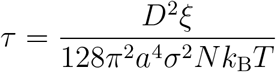

The viscosity in linear response regime is *η* = *P τ*^*R*^. Hence, the shear and normal stresses at linear response regime is obtained as follows,

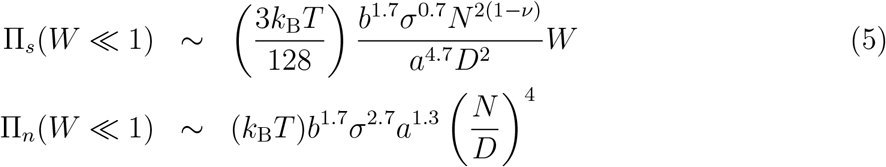

Now, I utilize the scaling argument to calculate the physical quantities at non-linear response regime. When the shear rate exceeds the critical shear rate 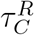, the shear extension must scale as *N*, similar to what the normal extension was in the equilibrium. This way, we determine the scaling function as Ξ = *W* ^1*/*5*ν*^ for the shear extension and Ξ = *W* ^*−*1*/*5*ν*^ for the normal extension. Doing so and some simplifications, we get the following similarity relations,

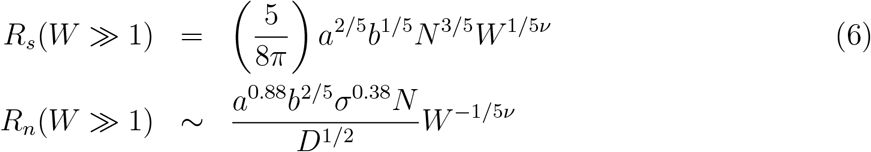

In the case of stresses, the scaling argument implies that the shear stress at non-linear regime, scales as *N* ^4^. This way, I determine the scaling function for stresses as Ξ = *W* ^(2*ν*+2)*/*(2*ν*+1)^. After doing simplifications, the following similarity relations obtained,

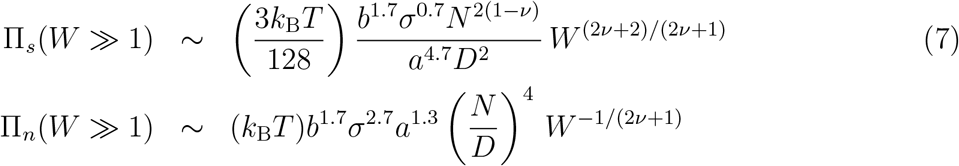

And finally, the kinetic friction coefficient at non-linear response regime is calculated by dividing shear stress by normal sterss,

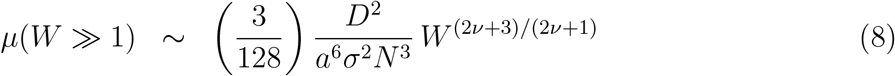

This is a very important result that we will see in the next sections it will be confirmed by MD simulations results.

### High shear rate scenario

At high shear rate, the chains continue stretching in the shear direction up to a moment when the top and bottom brush do not interpenetrate anymore. This is the moment when the normal extension equals half the wall distance. It means that when *R*_*n*_ = *D/*2 which leads to the following shear rate,

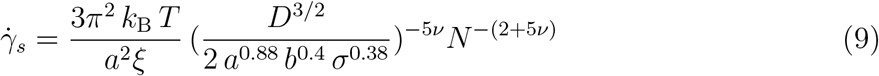

At 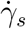 the equation of state by which we had build our scaling theory is changed. This is completely different scenario with respect to previous one. So we need to do everything from the scratch. The equation of state in this condition is the equation of state of a single brush which is given as follows,

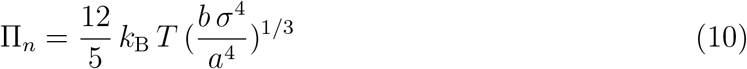

The equation above is calculated by using the grand potential of a single brush 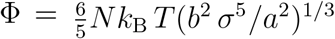, the Gibbs-Duhem relation Φ = *−*Π_*n*_ *V* and the normal extension of a single brush *R*_*n*_ = (*a*^2^ *b σ*)^1*/*3^*N/*2. On the other hand, the mean free path in this conditions is calculated as follows,

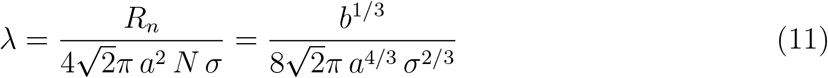

The mean free time is then calculated as follows,

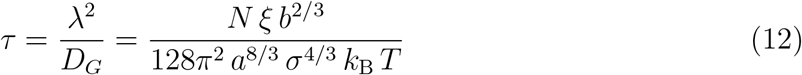

So we can now calculate the viscosity in this scenario as follows,

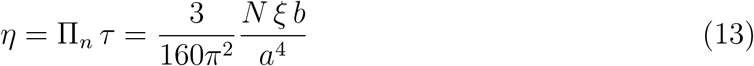

Now, we can use the scaling theory to check shear stress at high shear rates in this scenario. As before, we build an scaling function Ξ(*W*) = *W*^*α*^ and try to calculate *α* by considering the power of *N* . The phenomenological argument here is that when the chains are stretched in the shear direction, the shear stress must be proportional to *N* . This leads us to the equation 1 + (2*ν* + 1)*α* = 1 which gives us *α* = 0. This tells us that in the no-interpenetration scenario the shear stress is given as following,

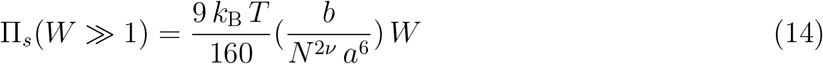

The normal stress is obtained by using the fact that it must become proportional to *N* ^3^ at high shear rate. This leads us to the equation 4 + (2*ν* + 1)*α* = 3 which clearly gives us *α* = *−*1*/*(2*ν* + 1). Therefore, the normal stress in this scenario is given as follows,

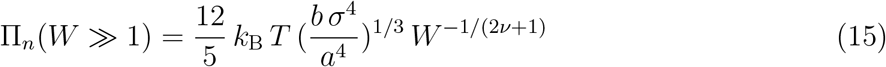

Therefore, the kinetic friction coefficient in this scenario becomes the following relation,

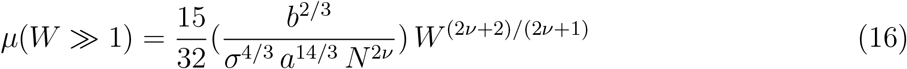

Table (**??**), summarizes universal power laws in nonlinear response regime.

### Molecular dynamic simulation

The molecular dynamic simulation is based upon calculation of trajectories of each particle in a discrete manner. The trajectories of each particle is calculated from the Newton’s equation of motion. In the present research, I use the Leap-Frog algorithm to discrete the equations of motion of monomers. This scheme calculates trajectories of monomers with a precision of the order of magnitude Δ *t*^4^ which is very precise. It also allows for small time step and faster calculations. In the simulations, the system is composed of two surfaces located at *z* = 0 and *z* = *D*. The surfaces are build by arranging atoms in a 2D simple cubic lattice. The area of the surfaces is *L*_*x*_ = 10*σ* times *L*_*y*_ = 20*σ*. The linear polymer chains of degree *N* are grafted to randomly chosen wall atoms. The periodic boundary conditions (PBC) is employed to mimic the bulk properties in the system. The shear motion is produced by moving the bottom wall atoms in the +*y* direction and the top wall atoms in the *−y* direction with speed *v/*2. The Lennard-Jones potential which make the hard-core excluded volume interactions between particles is given as follows,

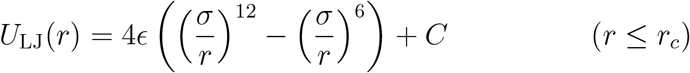

And the finitely-extensible-nonlinear-elastic (FENE) potential connects the monomers in the chains is given as follows,

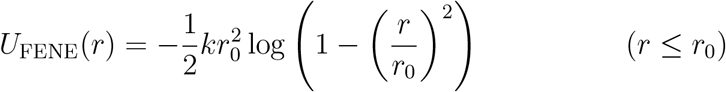

The Langevin thermostat is given as follows,

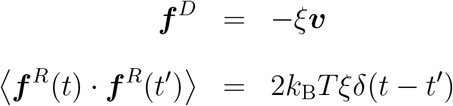

In the coarse-grained simulation all quantum mechanical degrees of freedom are integrated. It is called Kremer-Grest model.^21,22^ The simulations have run for 10^5^ time steps and to capture the most accurate results. Table (1), shows the numerical values of simulation parameters in my simulation.

**Table 1:** Numerical values of the simulation parameters.

| Quantity | Value | Quantity | Value | Quantity | Value | Quantity | Value |
| --- | --- | --- | --- | --- | --- | --- | --- |
| $dt$ | $2 \times 10^{-3}$ | $T$ | 1.68 | $r_c^{LJ}$ | $2^{1/6}$ | $b$ | 2.09 |
| $N$ | 30 | $\epsilon$ | 1 | $r_0$ | 1.5 | $\eta_s$ | 5 |
| $\sigma_g$ | $10^{-1}$ | $\sigma$ | 1 | $k$ | 30 | $D$ | 15 |
| $\xi$ | 5 | $a$ | 1 | $r_c^{DPD}$ | 2.3 | $\nu$ | 0.588 |

**Figure 1:**
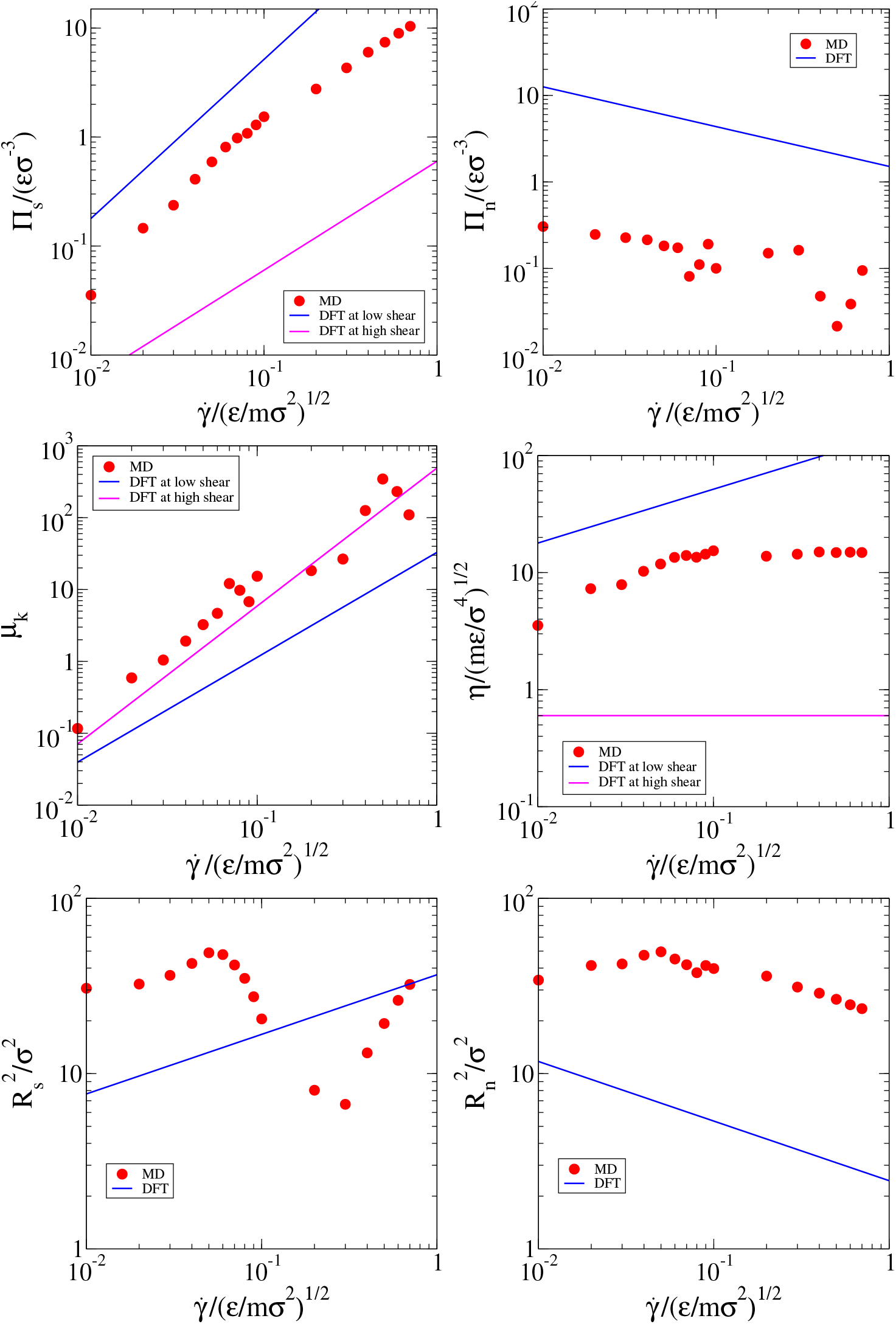
The shear and perpendicular stresses, the kinetic friction coefficient, the viscosity, the shear and perpendicular chain extensions as functions of the shear rate for MD and DFT calculations.

## Results and conclusions

The problem of the PBBs under stationary shear motion is studied by means of the DFT framework, the scaling theory and the MD simulations. Without any complicated calculations or extensive numerical simulations, one immediately infers that when two opposing polymer brushes intermediately compressed such that their chains interpenetrate into each other, the friction between them is larger than simple fluids. The reason is clear the interpenetrated chains produce friction. However, to find out the power laws we must use theoretical tools. So, my theoretical approach predicts that when two interpenetrating brushes are sheared, the shear stress scales as 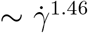, of course, as long as the brushes interpenetrate. This universal power law is very important because it correctly shows that if we use the equation of state of the polymer brush bilayer with interpenetrating chains, and apply the scaling theory to this equation of state, we get the correct power law. This is really amazing because molecular dynamic simulations confirm this power law well. The second scenario takes place when the chains stretch drastically in the shear direction such that eventually the normal chain extension reaches *D/*2 and at this moment the interpenetration zone vanishes. At this moment, the equation of state is changed and we have to apply the scaling theory to the equation of state of a single brush or non-interpenetrating brushes which are equal. When we do the same procedure on this equation of state, we surprisingly get the power law 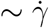 for the shear stress. This is really really amazing because both theory and simulations strongly agree on this scenario. In conclusion, the interpenetration between top and bottom brushes play the key role in the behavior of the system. If we take into account the explicit solvent molecules, it is always and in any conditions the solvent molecules concentrate in the middle of brushes and push the brush monomers out of the middle zone. It means that with explicit solvent molecules we never ever have an interpenetration zone between brushes. Note that you can not reach any interpenetration by compressing the brushes over each other because the solvents are always there. Therefore, with explicit solvents, the shear stress always scales linearly with the shear rate. Therefore, at the end of this article I would say that the works already published^5,13–15^ are completely wrong.

## Acknowledgments

I am honored to acknowledge the Cold Spring Harbor Laboratory (CSHL) and the Regeneron Pharmaceuticals Inc. for giving credit to my research. I also, thank Weizmann institute for the XMGrace software, the Wolfram Research for the Mathematica software and the GNU Inc. for the GFortran without which my research was not come into reality.

